# FeSBCP Analogue from Cyanobacteria: Insights from *in vitro* and *in silico* Studies

**DOI:** 10.1101/2025.08.30.673201

**Authors:** Aparna Boral, Titir De, Anwesha Banerjee, Tuyan Biswas, Bidisha Chakraborty, Devrani Mitra

## Abstract

The blue light responsive Cryptochrome/Photolyase Family harbours two important photoactivatable proteins - cryptochromes and photolyases. While cryptochromes are essentially photoreceptors with diverse biological activities including circadian rhythm, photolyases repair UV damaged cyclobutane pyrimidine dimers (CPD) or (6-4) pyrimidine-pyrimidone photoadducts. Both cryptochromes and photolyases share a common Photolyase Homology Region (PHR) where they harbour a light harvesting antenna chromophore and redox-active flavin di-nucleotide (FAD). Even though photolyases do not possess an extension in the C-terminal region unlike cryptochromes, a recent discovery of FeS cluster containing bacterial cryptochrome-photolyase proteins (FeSBCPs) present an interesting retreat from the conventional photolyase architecture. They possess an additional modular redox center, [4Fe4S] iron-sulfur cluster connected by conserved cysteine residues. Herein, we characterize a cyanobacterial FeSBCP from *Cyanobium sp*. using tools from bioinformatics, biochemistry and biophysics. Sequence analysis reveals substitution in the well-conserved aromatic residues, meant for electron transfer. *In silico* modeling and docking supported by electrophoretic mobility shift assay as well as spectroscopic measurements do suggest efficient binding and repair of CyPhrB to damaged single/double-stranded substrates containing 6-4 photolesion. Considering FeS clusters’ established roles in DNA binding/repair activities, we speculate the role of FeS clusters in FeSBCPs and existence of FeS-FAD-DNA triangle towards efficient electron transfer.

## Introduction

Sunlight is the ultimate source of energy used by life on earth. Genotoxic effects in living organisms are caused by detrimental DNA damage by exposure to the ultraviolet (UV) range of sunlight (Emmerich et al, 2020). Evolution has engendered robust DNA repair enzymes for repairing incorrect sequences or damages in DNA strands, thereby safeguarding the information encoded by them. DNA repair mechanisms can be for single-stranded (ss) DNA damage repair, direct reversal repair, nucleotide excision repair, base excision repair, and mismatch repair. For double-stranded (ds) DNA damage repair, homologous recombination and non-homologous end joining repairs are prominent. Two distinctive classes of photoactivatable proteins, the photolyases and cryptochromes, have evolved to correct UV radiation-induced DNA damage via direct reversal repair. These proteins are present in all three kingdoms of life (Geisselbrecht et al, 2012).

Cryptochrome/photolyase family (CPF) comprising of cryptochromes and photolyases are blue light (BL) sensitive photoactivatable proteins (Lin, C. & Todo, T. 2005). It has been observed that when DNA strands with adjacent pyrimidines (mostly thymines, TT) are exposed to UV radiation, new bonds develop between them. The formation of C6-C6 and C5-C5 bonds lead to cyclo-butane pyrimidine dimers (CPD), while the formation of C6-C4 bonds results in (6-4) photoproducts. The CPD reaction product is stable, whereas the product of the C5-C6 bond with the C4 carboxyl or amino group rapidly rearranges into the (6-4) photoproduct. Thus, photolyases can be classified based on the type of photoproduct they can repair (Sancar, 2008). One sub-class known as (6-4) photolyases, which primarily repairs (6-4) pyrimidine-pyrimidone photoproduct (6-4 photoproduct), was initially thought to be present only in eukaryotes, but was later also found in prokaryotes. The other type, known as CPD photolyases, repairs CPD and is present in all three kingdoms of life (Chaves et al, 2011) (Geisselbrecht et al, 2012). Photolyases (Phot) and their structurally-related counterparts, cryptochromes (CRY) photoreceptors, belong to the CRY/PHOT superfamily. Incidentally, both of them are activated by blue light owing to the presence of flavin di-nucleotide (FAD). While photolyases repair UV-induced TT lesion, cryptochromes predominantly regulate growth and development in plant systems and tune the circadian rhythm in animals and plants (Chaves et al, 2011) (Geisselbrecht et al, 2012) (Zhang et al, 2013).

Phylogenetically, all photolyases are thought to have diverged from a single ancestral CPD progenitor gene into Class I, Class II and Class III CPD photolyases. The (6-4) photolyases form another branch, believed to have evolved from Class I CPD photolyase. Single-strand specific DNA photolyases are known as CRY-DASH, where DASH stands for *<u>D</u>rosophila, <u>A</u>rabidopsis, <u>S</u>ynechocystis* and *<u>h</u>*uman (Kiontke et al., 2020). (6-4) photolyases can be further divided into classes based on the hierarchical groups they occur in, their structure and composition-Class I (eukaryotic), Class II (FeSBCP) and Class III (*Gloeobacter* (6-4) photolyase-like). Iron sulphur cluster-containing bacterial cryptochrome/photolyase (FeSBCP) subfamily has been discovered recently, but is evolutionarily most ancient. They are akin to (6-4) photolyases functionally but have significant differences based on sequence with all other subfamilies (Zhang et al, 2013). Thus, the group of FeSBCPs is placed at the largest phylogenetic distance from the other groups of the cryptochrome/photolyase family (Chaves et al. 2011).

Cryptochromes and photolyases share a similar structural makeup, known as Photolyase Homology Region (PHR). PHR has a bilobed architecture comprising two conserved domains which are connected by a variable loop. Photolyases usually exist as monomers, with their molecular masses ranging from 50 to 61 kDa (Kavakli et al. 2019). They possess two chromophores- a light harvesting cofactor and a catalytic FAD. The bilobed architecture is maintained in all structurally known class I DNA photolyases (photolyases primarily found in bacteria, fungi and plants) and plant cryptochromes as well as CRY-DASH with minor differences in the subdomains. The N-terminal α/β domain comprises five β-sheets and five α-helices, which forms the dinucleotide-binding domain housing the FAD cofactor. The FAD is lodged in U-shaped conformation of 14 interacting amino acid residues, majority of which are conserved. Besides the fully reduced and repair-active FADH-, two other redox states i.e. fully oxidized FAD_ox_ and semi-reduced FADH^o^ can be present in photolyases (Yamamoto et al., 2017). The latter two forms, via the process of photoactivation, get converted to the fully reduced version. The process of photoactivation is mediated by intra-protein electron transfer, where strategically placed tryptophan residues participate. The light harvesting cofactor, a nucleotide-like compound, is located in the cleft between the N and C terminal domains, approximately 17–18 Å away from the catalytic FAD chromophore. In some of the CRY/PHOT members, like in *E. coli*, the light harvester is MTHF (methenyltetrahydrofolate), while it is 8-hydroxy-7,8-didemethyl-5-deazariboflavin (8-HDF) in *Synechococcus elongatus* PhrB (SePhrB) (Chen et al, 2022), 6,7-dimethyl-8-ribityllumazine (DMRL) in *A. tumefaciens* FeSBCP, flavin mononucleotide (FMN) in *Thermus thermophilus* photolyase etc. (Kavakli et al. 2019).

Apart from the PHR, cryptochromes (except in CRY-DASH), possess a non-conserved Cryptochrome C-terminal Extension (CCE) domain. This domain is thought to have an indispensable role in functioning and regulation (Sarkar et al, 2025). The CCE domains are supposed to function like effector modules which get regulated by light. This folding or unfolding modifies their interaction with PHR and, thus, affects the entire conformation of the photoreceptors. The length of this domain greatly affects function (Yu et al, 2010). For example, in *Aradopsis thaliana* CRY1, the C terminal extension regulates plant morphogenesis and flowering under BL via COP1/SPA1 signaling (Sarkar et al, 2025). The FeSBCPs contain an additional Fe-S cluster in the C terminus alongside the other two redox active chromophores. Usually restricted to bacteria thriving in an iron-rich environment, the exact role of the Fe-S cluster in these proteins has not been wholly understood yet.

Despite FeSBCPs being abundant throughout the bacterial species, cyanobacteria have not been seen to possess FeSBCP, until SePhrB was reported recently (Chen et al., 2022). Cyanobacteria are well known to express CRY-DASH – the single strand-specific photolyases. In this manuscript, we investigate a new member of FeSBCP from another cyanobacterium, *Cyanobium* sp. (CyPhrB, (UNIPROT ID - B5IJS9)). This is the second report of FeSBCP from cyanobacteria to the best of our knowledge. Herein, we characterize CyPhrB using tools from structural bioinformatics and *in vitro* experiments. Following recombinant expression and purification, we identify the photolyase activity of CyPhrB. We further investigate the possibility of electron transfer to the UV-lesion site via multiple redox centers, including the Fe-S cluster. In this regard, we comb through multiple FeS clusters containing DNA repair enzymes and do a comparative analysis with CyPhrB.

## Materials and Methods

### *In silico* modelling and DNA-Protein Docking Analysis

Preliminary bioinformatics analysis reveals sequence homology of CyPhrB with other FeSBCP members, e.g. AtPhrB from *Agrobacterium tumifaciens* with known 6-4 photolyase activity. In absence of any experimental structure, we performed homology modelling of CyPhrB using MODELLER (Webb & Sali, 2016) using AtPhrB as template (PDB ID – 4DJA; Zhang et al., 2013). We then docked the modelled CyPhrB with undamaged DNA as well as repaired but flipped DNA. The latter was retrieved from the co-crystallised structure of *Drosophila melanogaster* (6-4) photolyase (PDB ID – 3CVY; Maul et al., 2008) using HDOCK webserver (Yan et al, 2017). We used visualisation tools like PyMOL and UCSF Chimera (Pettersen et al., 2004) not only to analyse potential docked conformations but also to perform sequence-structure analyses. We further used PDBePISA (Krissinel & Henrick, 2007) to understand the feasibility of modelled protein-small molecule (redox cofactors) as well as protein-DNA interactions.

### Phylogenetic Studies

Modelling studies reveal close sequence-structure relations of CyPhrB with other FeSBCP members with demonstrated (6-4) photolyase activity. We therefore performed a phylogeny analysis with 110 different 6-4 photolyases to understand the former’s evolutionary position. The phylogenetic tree was constructed using MEGAX software (Stetcher et al, 2020). Neighbour joining method with 1000 bootstrap replication values was used to make the analysis fast and error proof.

### Overexpression and Protein Purification for *in vitro* Studies

We transformed the codon optimized synthetic gene (GenScript) of CyPhrB cloned in pET28a into *Escherichia coli* C43 cells for protein overexpression. 4 litres of bacterial culture were grown in Luria Bertani Miller broth (HiMedia) at 37°C until O.D_600_ reached 0.6. We added 300 μM β-D-1 thiogalactopyranoside (IPTG) (SRL) to the culture for protein expression and kept it at 22°C overnight in the presence of blue light. After harvesting *E. coli* cells, the cell pellet was dissolved in a lysis buffer [Buffer A: 50 mM Tris, pH-8; 300 mM NaCl; 10% glycerol] with added lysozyme and protease inhibitor cocktail (Sigma). Cells were ruptured via sonication and supernatant with over-expressed CyPhrB (following SDS-PAGE analysis) was collected following high-speed centrifugation (Thermo Scientific). The supernatant fraction was then loaded onto the Ni-NTA affinity column (Qiagen), pre-equilibrated with buffer A. Non-specifically bound proteins were washed off the column using buffer B [50 mM Tris, pH-8; 300 mM NaCl; 10 mM Imidazole, 10% glycerol]. The CyPhrB fractions were eluted subsequently with buffer C [50 mM Tris, pH-8; 300 mM NaCl; 250 mM Imidazole, 10% glycerol]. CyPhrB containing fractions were pooled together, concentrated and subjected to size exclusion chromatography, using buffer D [50mM Tris, pH-8; 50 mM NaCl; 10% glycerol] - equilibrated ENrich SEC 70 10 column (Bio-Rad). Eluted fractions were tested for the presence of CyPhrB following resolution at 12-15% SDS-PAGE (Bio-Rad). Desired fractions were pooled, concentrated (Amicon, Millipore) and kept at -20°C for further use. Considering low protein yield, desalting (PD 10 desalting column, Sigma) was sometimes followed bypassing size exclusion chromatography for optimization of assay conditions and experiment replicates.

### UV-Vis Spectroscopy

We checked the purified fractions of CyPhrB by UV-Vis spectroscopy not only to understand the incorporation of redox cofactors but also to measure protein concentration (280nm, Molar Extinction Coefficient - 110600 M^-1^cm^-1^) for subsequent biochemical assay.

### DNA Damage and Repair Studies

We first considered the lyophilized ssDNA substrate (5’ACAGCGGTTGCAGGT3’) and dissolved it in nuclease free water. After dividing it into two batches, we measured the concentration using O.D_260_ value at the UV-Vis spectrophotometer. We then purged Argon or Nitrogen gas to both the batches for 20-30 minutes. One batch was kept in the dark for 3 hours and the other was exposed to UV-C (254nm, Philips) radiation for 3 hours. Fluorescence study was performed next to check for damage using undamaged DNA as control. Purified CyPhrB protein was then incubated with both the damaged and undamaged DNA. The mixtures were then exposed to BL for 1 hour at 4n followed by fluorescence measurement. Besides ssDNA damage and repair, we repeated fluorescence measurements with dsDNA as well. We dissolved the lyophilized (ss) complementary oligos in nuclease free water. 10μM of each oligo was annealed after heating at 95°C and successive overnight cooling in annealing buffer (10 mM Tris, pH 8.0; 20 mM NaCl). The dsDNA oligonucleotide was then set for damage and repair using the same protocol as that of the ssDNA strands.

### Electrophoretic Mobility Shift Assay (EMSA) for DNA Binding Study

We conducted EMSA to check CyPhrB’s affinity towards both undamaged as well as damaged ssDNA (5’ACAGCGGTTGCAGGT3’) and dsDNA. We added serially diluted CyPhrB (25µM – 0.09µM) to the assay mixture followed by 45 min incubation on ice. Protein-DNA complexes along with the control set (free DNA) were resolved in 10% native polyacrylamide gel for 1 hour at 150 V. We stained the gel using Sybr Gold in a 0.5X TBE buffer. The gel images were taken using the Gel Documentation System (Bio-Rad). We repeated the experiment for better resolution of the gels as well as to assure the consistency in results.

## Results

### CyPhrB: structure, sequence and evolution

The modelling of CyPhrB was done using MODELLER software (**FIG 1**). The best scoring modelled structure of CyPhrB does predict the presence of the FeS cluster. AtPhrB (*Agrobacterium tumefaciens* PhrB, ID – 4DJA), the closest structural and sequential homolog of CyPhrB and a member of FeSBCP family, is reported to possess four cysteine residues i.e., Cys350, Cys438, Cys441 and Cys454 (Zhang et al, 2013). These cysteine residues are essential for the FeS cluster binding. Indeed, multiple sequence alignment as well as CyPhrB do reveal that CyPhrB does possess those conserved cysteine residues (Cys359, Cys447, Cys450 and Cys463) (**FIG S1**) thus confirming the presence of [4Fe-4S] cluster. Further, the detailed interaction analysis of the FAD binding site, the antenna chromophore binding site and the FeS binding site of the modelled CyPhrB is shown in **Table S1**. This analysis and a very low R.M.S.D value (0.9Å) when compared with the AlphaFold predicted structure proves the authentication of our modelled structure. Functionally indispensable residues like His366, Tyr424, Tyr430 which interact with both FAD and DNA in AtPhrB and electron transmitting residues like Trp390 in AtPhrB, are also conserved in CyPhrB, suggesting the latter’s DNA reparative role. Such similarities inspired us to find out the evolutionary position of CyPhrB. The phylogenetic tree was thence constructed taking CyPhrB and 110 other (6-4) photolyases including AtPhrB as well as RsCryB (*Rhodobacter sphaeroides* Cryptochrome B, PDB ID - 3ZXS; Geisselbrecht et al., 2012). CyPhrB is found to assemble with the halobacter/cyanobacter (HC) group (**FIG 2A**). Its position is justified as the source organism of CyPhrB is a cyanobacterium. AtPhrB and RsCryB, albeit close homologs, are grouped under alphaproteobacteria group due to their origin. Notably, CyPhrB positions itself alongside another cyanobacterial FeSBCP, SePhrB from *Synechococcus elongatus* (PDB ID - 8IJY). Moreover, CyPhrB clearly falls under the FeSBCP group when its phylogenetic position was estimated taking plant and animal cryptochromes, photolyases, CRY-DASH and other FeSBCPs into consideration (**FIG S2**).

**Figure 1.**
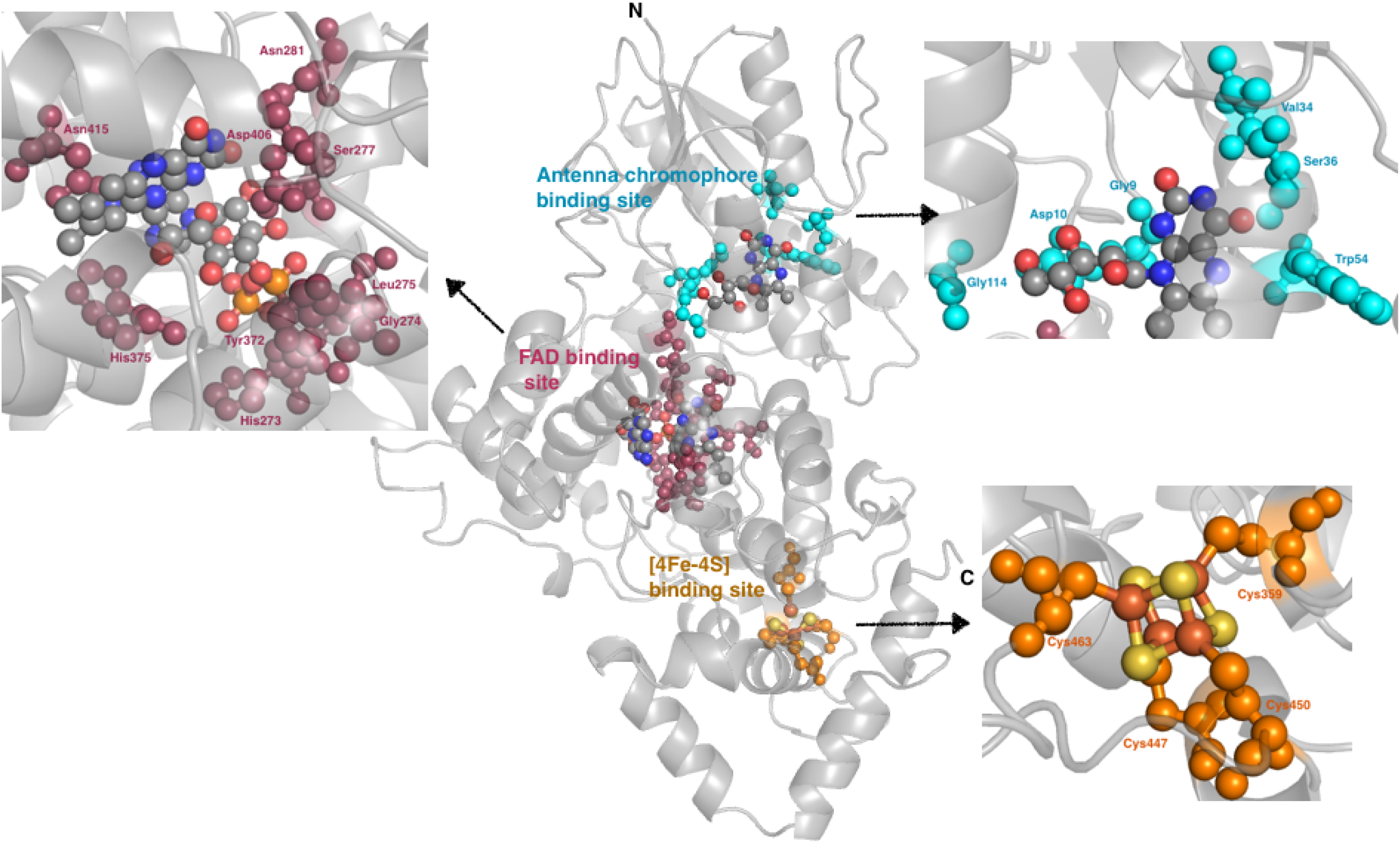
Structural analysis of CyPhrB. The modelled structure of CyPhrB is built using Modeller software. The Antenna chromophore binding site, the FAD binding site and the FeS cluster binding sites are magnified and shown in insets.

**Figure 2.**
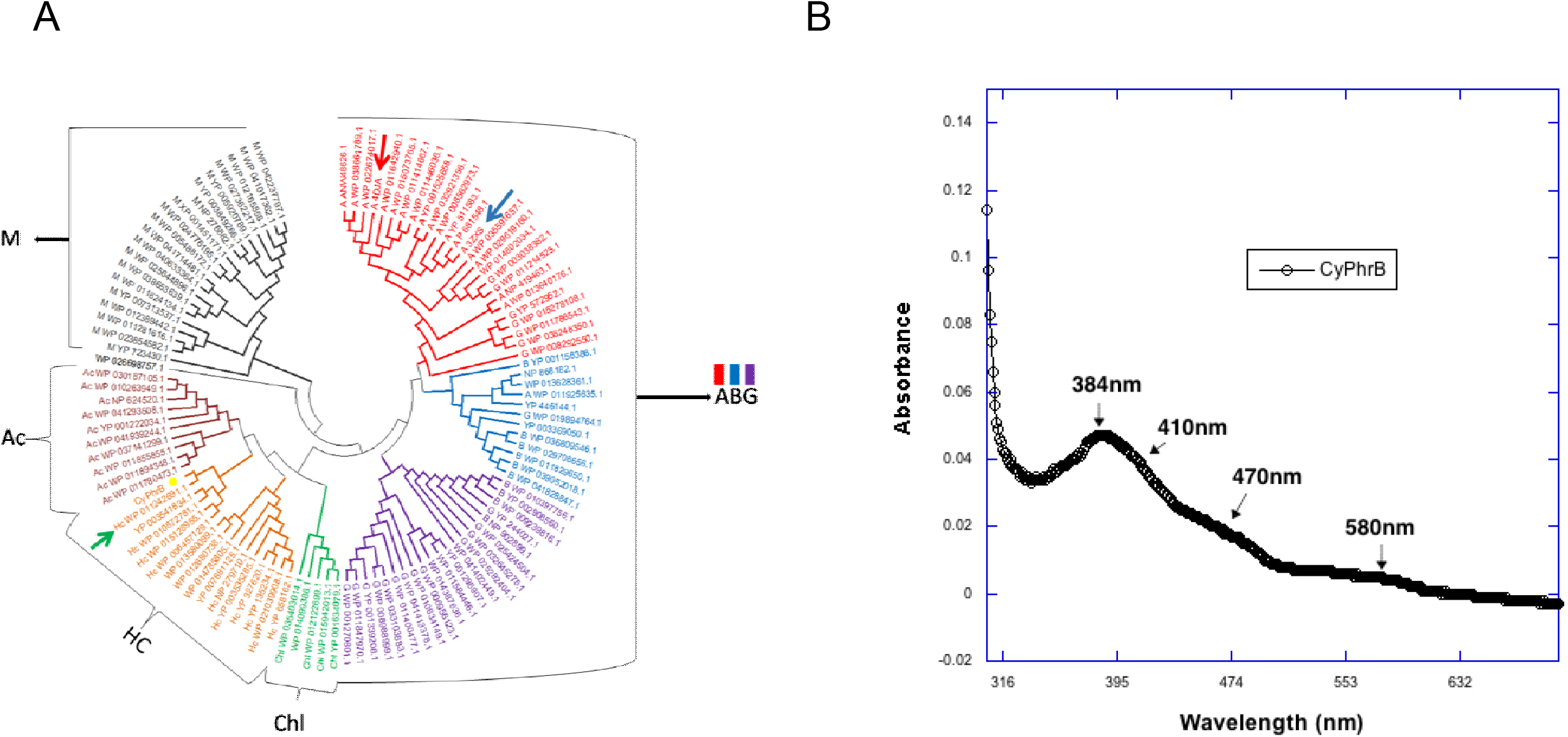
(A) **Phylogenetic analysis of CyPhrB**. The 111 prokaryotic (6-4) photolyases are divided into seven subgroups A,B,G (Alphaproteobacteria, Betaproteobacteria and Gammaproteobacteria), Chl (Chlorobi and Chloroflexi), HC (Halobacteria and Cyanobacteria), Ac (Actinobacteria and Acidobacteria), and M (Miscellaneous). The position of CyPhrB is shown by a yellow dot. The information of 110 sequences of 6-4 photolyases were taken from the work of Chen et al., 2022, the group names are given according to them. The sequence of CyPhrB was added to see its phylogenetic position. The position of AtPhrB is shown by a red arrow, RsCryB is shown by blue arrow and SePhrB is shown by green arrow. (B) **The UV-Vis spectrum of CyPhrB**. The important features are shown by arrows. The peak at 384 nm is assigned to the presence of antenna chromophore, the absorbance at 410 nm signifies the presence of FeS cluster, the absorbance at 470 nm and 580 nm marks the presence of semi reduced FAD.

SDS-PAGE analysis reveals the molecular weight of CyPhrB to be around 60kDa (**FIG S3**). The UV-Vis spectrum detects absorbance between 400- 420 nm. The shoulder at 410 nm is possibly due to the presence of a FeS cluster in CyPhrB, similar to that in SePhrB (Chen et al, 2022). The peaks obtained at 384 nm, 470 nm and 580 nm are possibly due to the presence of antenna chromophore and semi reduced FAD respectively **(FIG 2B**). Initially, since the template for modelling CyPhrB was AtPhrB, the choice of antenna chromophore was 6,7-dimethyl-8-ribityllumazine (DLZ) for obvious reasons. When residues of the DLZ binding site were compared with that of the DLZ binding site of AtPhrB, it is seen that it has approximately 43% residue identity. We also tried with other antenna chromophores available like 8-HDF (reported in SePhrB) and MTHF (reported in *Arabidopsis thaliana* Cry3). Residue level analyses reveal approximately 45% residue identity with that of the residues interacting with 8-HDF in SePhrB, but it does not show any identity with the MTHF interacting residues of *Arabidopsis thaliana* Cry3. While the antenna chromophore 8-HDF is not synthesized by *E*.*coli* (Chen et al, 2022), DLZ and MTHF are readily synthesized. In absence of 8-HDF, CyPhrB might have incorporated DLZ as an antenna chromophore with a resultant 384 nm absorption similar to AtPhrB (Oberpichler et al., 2011)

### CyPhrB : A cyanobacterial (6-4) photolyase

In terms of function, FeBCPs are assumed to resemble (6-4) photolyases. A histidine residue (His366 of AtPhrB) is conserved across both FeSBCPs and photolyases and essential for DNA repair. The homologous His375 in CyPhrB (**FIG S1**) does exhibit hydrogen bonding with the TT dimer in the modelled docked structure (**FIG S1 and TABLE S1**). Apart from residue-level homology, there are certain differences as well. For example, a certain histidine (His369 in *Drosophila melanogaster* 6-4 photolyase) is conserved in all 6-4 photolyases except in FeSBCPs. In the latter, His is replaced with leucine, e.g. Leu379 in CyPhrB (**FIG S1**). Critical intra-FeSBCP differences include - a certain methionine residue (Met399 in SePhrB, a cyanobacterial FeSBCP) is present near the N5 position of the flavin chromophore. Similarly, CyPhrB being a cyanobacterial FeSBCP too has a methionine in the homologous position (Met 412). However, the bacterial FeSBCPs like AtPhrB and RsCryB possess a glutamate residue (**FIG S1**) instead. Again, besides the similarities between the cyanobacterial FeSBCPs, perhaps the most striking difference between CyPhrB and SePhrB is the substitution of Trp338 (homologous to Trp342 in AtPhrB) with glutamate (Glu350). The only conserved Trp residue in the conventional Trp-triad is Trp399, homologous to Trp390 in AtPhrB. Incidentally, Trp342 and Trp390 are found to be essential in AtPhrB for electron transfer process (Holub et al., 2018) (**FIG S1**). Another major difference lies in Phe400, which is Tyr391 in *At*PhrB. Interestingly, both the cyanobacterial PhrBs have phenylalanine instead of tryptophan at this critical position. However, as per the results of Holub et al., 2018, the substitution of Tyr391 with phenylalanine should not be a hindrance to photoreduction.

Next, our focus was to understand the binding affinity of CyPhrB with the kinked DNA and for that, we performed *in silico* docking. The docking of CyPhrB was performed using HDOCK webserver and the best scoring structure was taken for further analyses. The substrate DNA used for docking was taken from the co crystallised structure of *Drosophila melanogaster* 6-4 photolyase (PDB ID - 3CVY). When compared with AtPhrB, RsCryB and SePhrB, the overall DNA binding regions of all the docked structures are seen to be quite similar (**FIG 3 and FIG S1**). Also the protein-DNA interaction analyses conducted using PDBePISA does reveal that the DNA binding site and the residues are indeed common among the FeSBCPs (**Table S1**). Further, it is seen that CyPhrB binds better with the flipped TT dimer containing DNA than with its undamaged counterpart, in agreement with typical photolyase activity.

**Figure 3.**
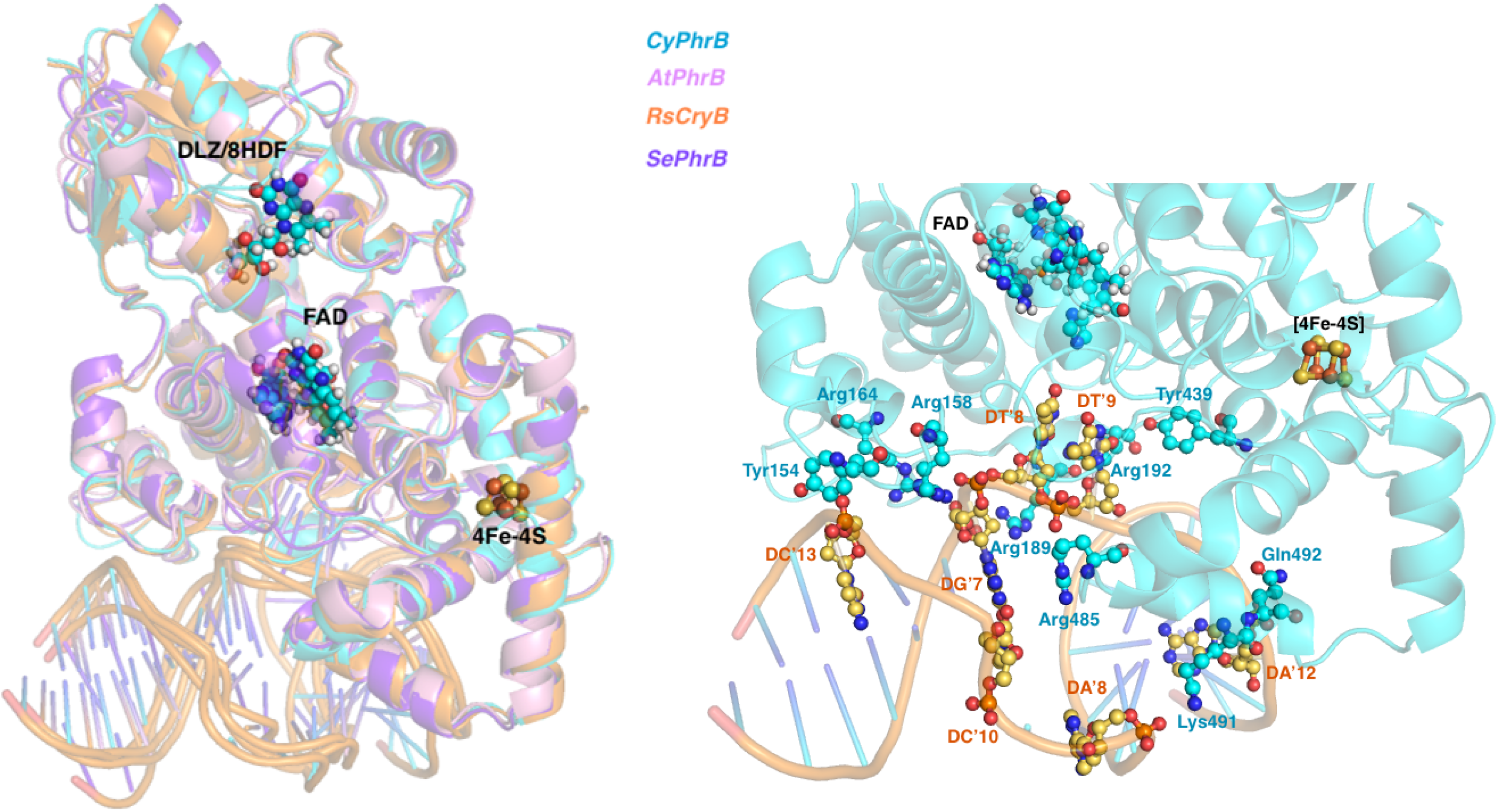
DNA binding analysis of CyPhrB by *in silico* docking. CyPhrB (model) is docked with the substrate DNA. The best scoring docked structure is then compared with the best scoring docked structures of AtPhrB, RsCryB and SePhrB. The substrate DNA for all the docked structures are obtained from the co crystallised structure of Drosophila melanogaster 6-4 photolyase (PDB ID – 3CVY). The magnified image of the DNA binding site along with the residues involved in protein-DNA interaction of CyPhrB is also shown.

The above-mentioned confirmation made us look more into the DNA repair property of CyPhrB. Initially, an aqueous solution of the lyophilised ss- as well as dsDNA oligonucleotides (DNA substrate sequences obtained PDB ID – 3CVY) was prepared and the O.D_260_ was kept at ∼0.5. Then one of each set of ss- and dsDNA were kept under UV-C radiation for 3 hours and another set of ss and ds DNA were kept in dark as a negative control. The UV exposed as well as the control sets of ss- and dsDNAs were then checked in fluorescence spectroscopy. Different parameters i.e., excitation wavelength, emission wavelength range, slit size etc. were carefully calibrated. The strong Raman peak of water was also excluded so as to reduce the ambiguity of the result. After finalizing the parameters, we excited the DNA oligonucleotides at 260 nm and the emission spectrum was taken in the range of 280-500 nm, considering the appearance of Raman peak beyond 500 nm. An aqueous solution of UV-irradiated thymine exhibits an increase in the fluorescence intensity around 375 nm when excited at 260 nm (Nagpal et al, 2021) – the authors assigned this peak as the formation of the 6-4 photo lesion. In our case, when compared with the control set, the UV irradiated set of both ss- and dsDNAs do show an increase in fluorescence intensity around 355nm (**FIG 4A, C**). The DNA sets used here are oligonucleotides – therefore, a blue shift in the peak is possible due to the hydrophobic environment. Moreover, it is observed that the increase in fluorescence intensity is more pronounced in the ssDNA than in the dsDNA. This might be due to the fact that in case of ss DNA, the TT dimer is more exposed for fluorescence signalling when compared to the dsDNA.

**Figure 4.**
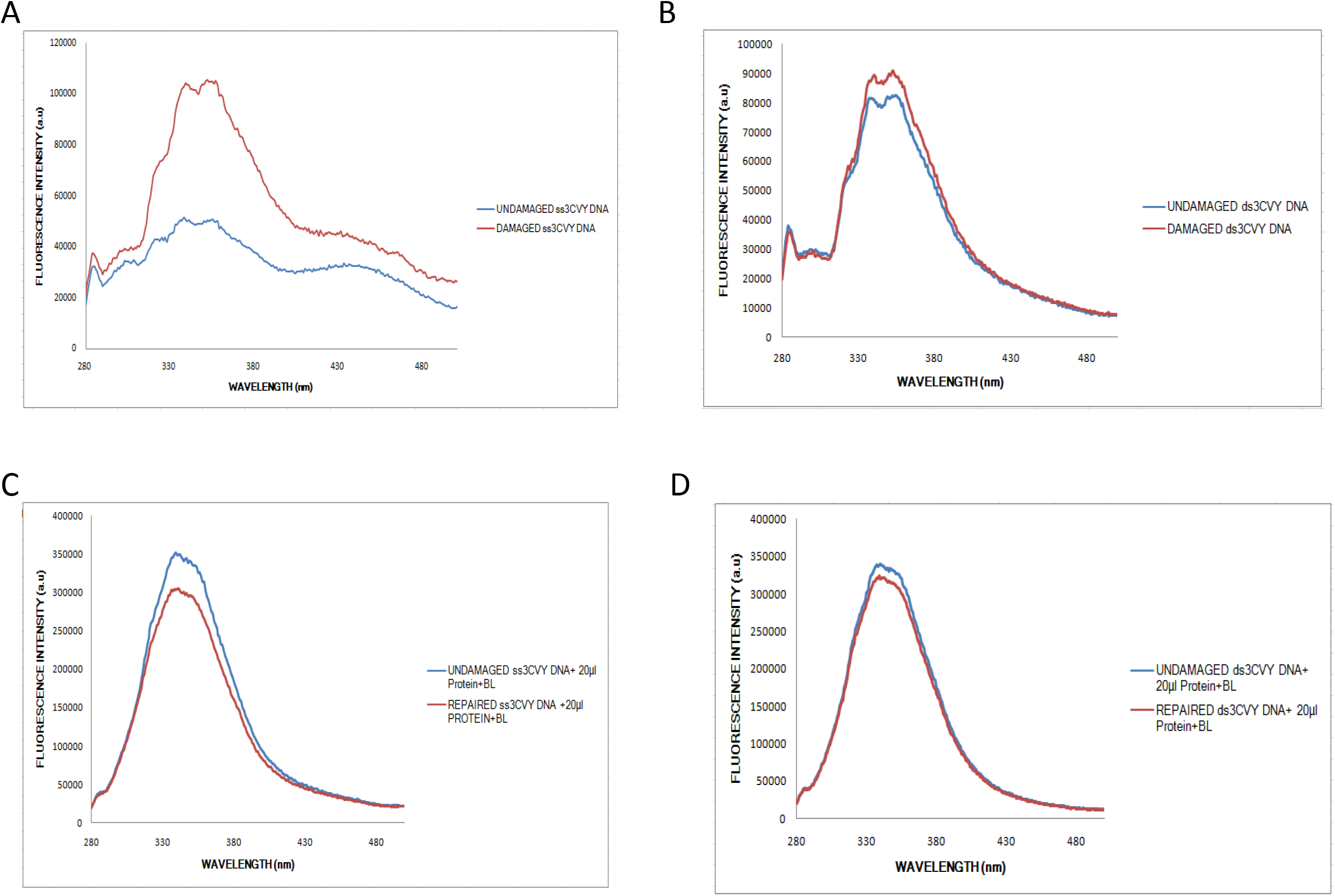
Fluorescence emission spectra of single (ss) as well as double stranded (ds) DNA substrates with (damaged) and without (6-4) photolesion (undamaged). (A and C) – Damaged vs. Undamaged ss- and dsDNA. (B and D) – Repaired vs. Undamaged ss- and dsDNA in presence of CyPhrB.

For the repair assay, we incubated the homogeneously purified CyPhrB with the undamaged and damaged ss- as well as dsDNA oligonucleotides in presence of BL for 1 hour. We then measured fluorescence for any consequent changes. Interestingly, both the undamaged and the damaged-turned-repaired ss- as well as dsDNAs showed almost the same spectrum. The damaged-turned-repaired DNA’s spectrum is in fact even slightly of lesser intensity than its undamaged counterpart (**FIG 4B, D**). The pattern of emission spectra is indicative of repairment of the damaged ss/dsDNA upon CyPhrB incubation in the presence of BL. The dramatic increase in the fluorescence intensity in both damaged and undamaged DNA following the addition of protein is due to the overlapping intrinsic fluorescence properties of the aromatic amino acids present in the protein (Yang et al, 2015).

We next conducted EMSA to validate results obtained from *in silico* docking as well as fluorescence spectroscopy. Our EMSA results clearly show better binding affinity with the damaged ss- and dsDNA oligonucleotides than their undamaged counterparts where the K_d_ of the damaged ss and dsDNAs ranges from 400-300 nM (**FIG 5**).

**Figure 5.**
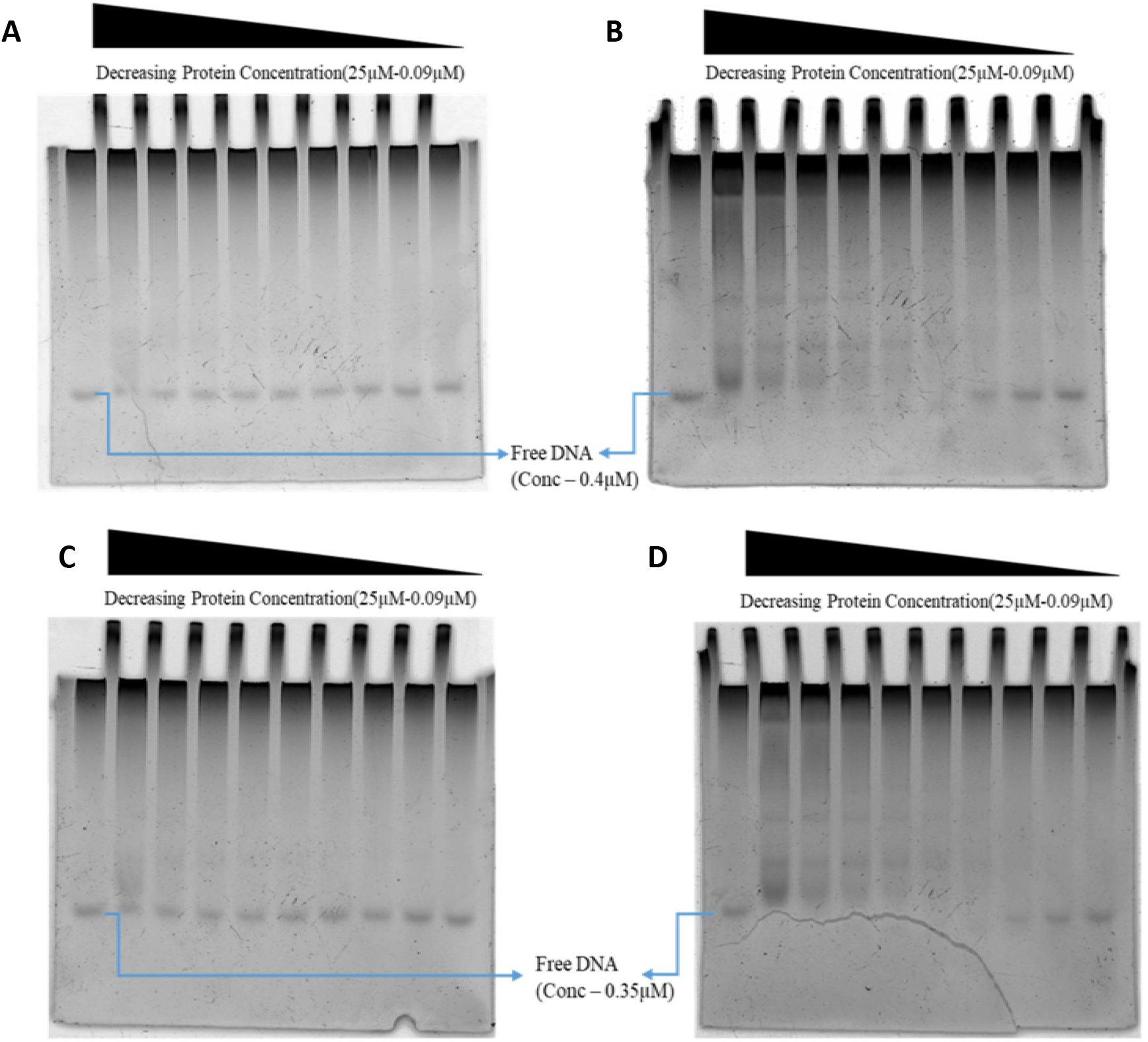
Electrophoretic Mobility Shift Assay (EMSA) of CyPhrB with undamaged and damaged DNA substrates. The undamaged substrate sequences were considered from Drosophila (6-4) photolyase’s substrate as referred in the PDB ID 3CVY. CyPhrB was made to interact with (A) ss undamaged DNA, (B) ss damaged DNA, (C) ds undamaged DNA, (D) ds damaged DNA, in presence of blue light.

### CyPhrB: the role of the FeS cluster

Although FeSBCPs are very common across all bacterial species, FeSBCPs from cyanobacteria are not studied barring one (SePhrB). CyPhrB is mostly possibly the second cyanobacterial FeSBCP that we investigate herein. The CyPhrB protein was docked with the Dm6-4Phot’s substrate DNA whose TT dimer was protruded out of the dsDNA. The docked structures of CyPhrB, SePhrB, AtPhrB, and RsCyrB with the best scores, and the co-crystal structure of Dm6-4Phot were taken for further analysis. We then computed the protein-DNA interface area of the docked CyPhrB structure and compared it with the co-crystal structure of Dm6-4Phot. The value of the gain in solvation free energy was also checked and compared to understand whether the binding of DNA to CyPhrB is spontaneous or not. As seen from **Table S1**, the areas of the DNA-bound interfaces in both cases (*in silico* docked structure of CyPhrB and Dm6-4Phot) are highly comparable. Moreover, the negative value of gain in solvation energy upon complex formation indicates favourable substrate DNA binding in CyPhrB, similar to Dm6-4Phot. The interface areas, gains in solvation free energy as well as the number of hydrogen bonding/salt bridge interactions for all the redox co-factors (antenna, FAD and FeS cluster) of CyPhrB are distinctly similar to other PDB structures, e.g. AtPhrB - 4DJA, RsCryB -3ZXS and SePhrB - 8IJY [**Table S1**].

## Discussion

Unlike most cryptochromes, photolyases possess a light-regulated mechanism of DNA photoproduct repair. The antenna chromophore first absorbs the BL photon and then transfers it to FADH^-^ via fluorescence resonance energy transfer (FRET). Next, the FADH^-^ gets excited (FADH^-*^) and transfers an electron to the TT photolesion, which then gets repaired. Subsequently, the electron gets back to the flavin and its fully reduced (FADH^-^) resting state is restored (Aguida et al, 2024). Further, unlike cryptochromes, in photolyases, the catalytically active form of flavin is FADH^-^. FADH^-^ is the only redox state that can catalyze DNA repair. Absence of this state, results in loss of DNA repair activity of the protein. While fully reduced form is available *in vivo*, oxidized or semi-reduced forms are mostly notable *in vitro*, especially in photolyases purified under aerobic conditions (Holub et al., 2018). Nevertheless, photoreduction to FADH^-^ is one of the most important requirements for the DNA repair activity of the photolyases. In case of cryptochromes, a conserved Asp396 (*Arabidopsis thaliana* Cry1) protonates the anionic radical of FAD to form the active signalling FADH° radical state. All non-DNA repair members of CPF possess this conserved aspartate residue. This negatively charged residue provides the flavin its biologically active semi-reduced state in cryptochromes. In case of photolyases, neutral asparagine or positively charged lysine residue is present in the homologous position of the aspartate (Sarkar et al, 2025). This single residue alteration is extremely critical as it is the diverging point between the signalling proteins (cryptochromes) and DNA repair enzymes (photolyases) (Burney et al, 2012). Interestingly, a single mutation of Asp396 to Asn could render an additional DNA repair property to the *Arabidopsis thaliana* Cry1 (Burney et al, 2012). CyPhrB, indeed possesses an Asparagine (Asn415) in the homologous position and is therefore a DNA repair enzyme. Perhaps the most interesting observation in CyPhrB is the departure from the conventional Trp-triad or at least a triad consisting of three aromatic residues as observed in its nearest cyanobacterial homologue, SePhrB. CyPhrB possesses Glu350, Trp399 and Phe400 instead of the conventional ones meant for electron transfer. Aromatic amino acids, especially tryptophans, are most befitting for electron transfer mechanism due to their possession of aromatic side chains, low redox potential and ability to form stable radical intermediates for the deprotonated forms (Holub et al., 2018). It is possible that Glu350 stabilizes the neighbouring radical Trp state via lowering the free energy barrier through facilitating proton transfer. However, the precise role of this unique glutamate substitution remains to be validated in future experiments. Glutamate is actually known to facilitate proton coupled electron transfer (PCET) between two tyrosine residues in ribonucleotide reductase enzyme from *E. coli* (Reinhardt et al, 2021). Incidentally, PCET is essential for the repair of 6-4 photolesion as well. Three fundamental processes are necessary to achieve this phenomenon - perception of a BL photon, donation of an electron from the redox co-factor and a proton transfer from the active site histidine residue (CyPhrB’s His375 homologues). In fact, mutation of this critical histidine residue in (6-4) photolyase has completely abrogated repair activity (Zhong, 2015)..

One particular methionine (Met399 in SePhrB and Met412 in CyPhrB) is close to the N5 position of the flavin chromophore whereas the bacterial FeSBCPs have a glutamate in the homologous position (**FIG S1**). This is an interesting observation as methionine is a redox active residue and does donate an electron in some cases (Chen et al., 2022). For example, a Cys57Met mutation in *C. reinhardtii* LOV1 does show that the methionine can form the covalent adduct with the N5 of FMN upon BL irradiation, thus producing the biologically active reduced form of flavin. The methionine at the same position in SePhrB and CyPhrB too might have a role in flavin reduction and consequent DNA repair. Furthermore, a certain Leucine (Leu366 in SePhrB and Leu379 in CyPhrB) (**FIG S1**) though is not directly involved in DNA repair but when mutated, does lower DNA repair activity as it has an important role in FAD binding in FeSBCPs (Chen et al., 2022).

The CPF members share a conserved architecture in the chromophore binding domains. Two chromophores- a light harvesting antenna, most commonly methenyl-tetrahydrofolate (MTHF), is bound to the N-terminal α/β domain and the catalytic FAD is bound to the C-terminal helical domain. The conserved helices are arranged in a Rossmann fold around the FAD cofactor and form the DNA binding domain which is a conserved structure but show differences in amino acid composition across the superfamily (Lamparter et al., 2014). There are also C-terminal extensions (CTE or CCEs) which vary in length and are poorly conserved even among cryptochromes of the same species like in *Arabidopsis thaliana*-Cry1 and Cry2. Majority of the photolyases lack such C-terminal extensions (Chaves et al., 2011). These CTEs are believed to be pivotal in cryptochromes having a varied range of functions. In a sub-group of prokaryotic photolyases e.g. FeSBCPs, the CTE houses a redox active 4Fe-4S cluster having functions that have remained elusive till date. The iron is coordinated with four cysteine residues of which three lie in the variable CTE region. When compared with other FeS cluster containing DNA binding proteins, the 4Fe-4S cluster may putatively act as a clip in the C-terminal domain of the protein, act as a lesion detector, aid in DNA binding or enable reversible electron cycling from the bound lesion to the FAD co-factor (White & Dillingham 2012). As already proven via various studies, FeS clusters containing proteins are able to communicate and synchronise their activities by long range DNA mediated charge transfer (Barton et al., 2011). The FeS cluster in these photolyase subsets may also enable such communication since DNA photolyase and cryptochromes as well as few other DNA repair proteins are usually found in the neighbourhood as noticed in the STRING database (results not shown).

Iron-sulfur clusters, ‘nature’s modular, multipurpose structures’ are known for their redox abilities in prokaryotes as well as eukaryotes (Beinert et al., 1997). Albeit present in various forms, the common forms are 2Fe-2S, 3Fe-4S and 4Fe-4S. FeS clusters are generally found as cofactors in enzymes and they usually help in carrying out the Lewis acid-base reactions found in the active site of mitochondrial aconitase and radical S-adenosylmethionine (SAM) enzymes. They are also known to regulate gene expression (Mettert & Kiley, 2014) in response to oxidative stress, by their presence in superoxide responsive (SoxR) proteins; varying oxygen levels, by fumarate-nitrate reduction (FNR) proteins; and iron levels, via Iron Response Proteins including IRP1 and IRP2. FeS clusters do play critical roles in DNA replication and repair. These clusters also participate in the maintenance of the structural stability of enzymes as well as that of the DNA. Moreover, the role of FeS clusters in electron transport is the most well-established one. A simple, successive change of the overall cluster charge, e.g. from [4Fe-4S]^0^ to [4Fe-4S]^2+^, or [4Fe-4S]^2+^ to [4Fe-4S]^3+^, FeS clusters mediate catalysis in the Fe-protein of nitrogenase (Mitra et al., 2013) or, participate in DNA repair activity (Boal et al., 2005). These ‘modular, multipurpose’ clusters possess absolutely necessary roles in the mitochondrial electron transport chain (ETC), and are found in complexes I, II and III of the ETC (Read et al. 2021).

Charge transfer requires kinetic feasibility and donor-acceptor coupling with respect to optimum distance of transfer as well as complementary redox potentials. The work by (Teo et al. 2019), discusses one general, kinetically favourable model for DNA-mediated charge transfer between FeS cluster containing proteins, a cofactor which is found in a variety of domains, majorly in proteins associated with DNA repair and replication. The proteins DNA polymerase alpha and primer p58c were considered in the kinetic modelling which led to the conclusion that the FeS cluster seems to be kinetically favoured to donate electrons to the bound DNA duplex, and the duplex can donate electron holes, not electrons to the protein. The model also highlighted the importance of redox active amino acids, tyrosine, tryptophan and methionine in the electron-hole transfer. Transfer of hole or electron across the duplex can consist of electronic couplings that mainly involve guanine (for hole transfer) or thymine (for excess electron transfer).

Looking deeper into the chromophore lining residues we observed that few very important residues like His366, Tyr424 and Tyr430 that interact both with FAD and DNA present in AtPhrB are also conserved in CyPhrB. One of the most important electron transferring residues, Trp390 of AtPhrB is also conserved in CyPhrB. Since there are numerous structurally characterised FeS containing proteins, and quite a few of them are involved in DNA transaction processes, we tried gaining further insights about the still elusive role of FeS clusters in FeSBCPs by residue level comparison of the [4Fe-4S] cluster containing DNA binding proteins with the FeSBCPs. We considered - MutY (PDB ID – 1RRQ) (Fromme et al., 2004), which repairs inappropriately inserted adenine residues, as a result of 8-oxoG mutation; DinG (PDB ID - 6FWS) (Cheng & Wigley, 2018) which functions in the nucleotide excision repair and recombinational DNA repair pathways; DNA2 (PDB ID - 5EAN) (Zhou et al., 2015), which is a nuclease-helicase that processes DNA double strand breaks, Okazaki fragments and stalled replication forks; AddB (PDB ID -3U44) (Saikrishnan et al., 2012) which is a Chi sequence recognition protein therefore helping in DNA break repair. In these proteins, the FeS clusters either play the role of structure stabilisation, acting as a clip, and/or, the redox state of the cluster thereby regulating the protein activity. These proteins have redox active FeS clusters, as reported. Upon matching the clashing residues of these proteins with the bound DNA via UCSF Chimera it was found that all of them are abundant in arginine, serine and proline residues in their DNA binding domains, which are less than 3.4Å away from the DNA backbone. This is no exception for FeSBCPs as well, as noticed from their docked structures. The distance measurements in all cases revealed similar proximity of FeS clusters from the DNA binding sites. In CyPhrB, the distance is (**FIG S4**) 17.1Å. Further, the distances in MutY, DinG, Dna2 and AddAB are 12.8Å, 12.2Å, 9.8Å and 21.2Å respectively. Again, the FeS cluster is approximately 16.1Å away from the FAD cofactor in CyPhrB, while the FAD is 8.7Å away from the substrate DNA. Moreover, several redox active residues like Tyr446, Tyr439, Tyr433 and Trp318 line up the FeS cluster-FAD route in CyPhrB, making electron transfer feasible. Since the FeS clusters regulate electron cycling between DNA and the other cofactors, this can be extrapolated to the FeSBCPs too. The FeS cluster in CyPhrB may be involved in electron transfer between cofactors and DNA, having a role in lesion detection and lesion repair, as well as communication with other DNA repair proteins via DNA mediated charge transfer processes. We speculate that a triangular electron transfer chain between FAD, FeS cluster and bound DNA might exist in FeSBCPs, with FeS clusters bolstering damage detection and electron transfer processes. However, this possibility needs to be validated via future experiments.

## Conclusion

FeSBCPs though common in bacteria, this is the second report of a cyanobacterial FeSBCP to the best of our knowledge. CyPhrB is sequentially, structurally as well as evolutionarily close to other bacterial FeSBCPs. However, CyPhrB is more closely aligned with its cyanobacterial counterpart, SePhrB. Perhaps the most striking difference between bacterial and cyanobacterial FeSBCP is the substitution of electron-transferring tyrosine residue with phenylalanine (Phe400 in CyPhrB). Both bacterial and cyanobacterial FeSBCPs possess four conserved cysteine residues that harbour an additional [4Fe-4S] cluster at the CTE. Moreover, the DNA binding residues reported in AtPhrB are also conserved in CyPhrB. While both *in silico* docking and EMSA studies are suggestive of higher DNA binding affinity of CyPhrB with damaged ss/dsDNA over undamaged ones, fluorescence measurements indicate (6-4) photolesion repair activity. We further hypothesize that FeS – FAD – substrate DNA forms an approximate electron transfer triangle, thereby elucidating the possible role of dual redox sensors in FeSBCPs. Deviation from the conventional electron transfer triad along with the presence of nature’s one of the most efficient redox centers, makes future FeSBCPs research fascinating. It will be worth dissecting the precise electron transfer pathway that possibly exists along the arms of FeS-FAD-DNA triangle.

## SI Figure Legends

**Figure S1**. Multiple Sequence Alignment (MSA) of CyPhrB, AtPhrB, RsCryB and SePhrB sequences. The DNA binding residues are shown in boxes, the cysteine residues required for FeS binding are conserved and shown with solid circles and the electron transfer residues are shown with solid stars.

**Figure S2**. Phylogenetic analysis of CyPhrB. A comparative phylogenetic tree of plant and animal cryptochromes, CRY-DASH, FeSBCPs, CPF1, CPF2 along with DNA photolyases shows their evolution into distinct branches. The position of CyPhrB is depicted with a yellow dot.

**Figure S3**. SDS-PAGE image and corresponding UV-Vis spectra of gel filtered fractions of CyPhrB, showing purified protein fractions at 60 kDa.

**Figure S4**. Distance analysis between FeS and TT residues in FeS containing DNA repair enzymes. In centre distance analyses between FAD, TT dimer and FeS in CyPhrB.

## Acknowledgements

Devrani Mitra and Aparna Boral acknowledge funding from the Department of Science and Technology and Biotechnology, Govt. of West Bengal, India (2400 (Sanc.)/STBT-13015/6/2021-ST SEC). Aparna Boral further acknowledges Indian Council of Medical Research (45/06/2022-/BIO/BMS) for senior research fellowship. The authors acknowledge DST-FIST and DBT-BUILDER grants provided to the Central Instrumentation Facility, Department of Life Sciences, Presidency University. The authors sincerely thank the past and present lab members, especially Sk. Asik Ali, Dr. Anwesha Deb and Debarshi Bose for helpful discussions. The authors are also grateful to the Department of Chemistry, Presidency University, especially Profs. Arnab Halder and Gandhi Kr. Kar, for providing the instrumentation facility during the initial phases of fluorescence experiments.

## Conflict of interest

The authors declare no conflict of interest.

## Author contribution

**Aparna Boral** - Methodology, validation, formal analysis, investigation, data curation, writing (original draft), editing and visualization; **Titir De** - Methodology, validation, investigation, editing; **Anwesha Banerjee** - Writing, Formal analysis, investigation, editing and visualization; **Tuyan Biswas** - Methodology; **Bidisha Chakraborty** - Methodology, editing; **Devrani Mitra** - Conceptualization, Methodology, validation, formal analysis, investigation, data curation, writing (original draft), writing (review and editing), visualization, supervision, project administration and fund acquisition.

## Notes

### Competing Interest Statement

The authors have declared no competing interest.

