## Supplementary Information for "FeSBCP Analogue from Cyanobacteria: Insights from *in vitro* and *in silico* Studies"

#### Table S1

| | | | | | | | | | |
|---|---|---|---|---|---|---|---|---|---|
| **Name** | **Interface Area, (Å$^2$)** | **Gain in solvation energy upon interface formation, ΔG (kcal/mol)** | **Molecules forming complexes** | **Hydrogen bonds** | | | **Covalent bonds** | | |
| CyPhrB | 642.2 | -6.2 | PROTEIN+FAD [CHROMOPHORE, REDOX CENTRE] | **PROTEIN** | **DISTANCE Å** | **LIGAND** | **PROTEIN** | **DISTANCE Å** | **LIGAND** |
| | | | | A:GLY 274(N) | 3.64 | B:FAD 525(O1P) | | | |
| | | | | A:LEU 275(N) | 2.93 | B:FAD 525(O1P) | | | |
| | | | | A:TYR 372(OH) | 3.28 | B:FAD 525(O1P) | | | |
| | | | | A:SER 277(OG) | 3.32 | B:FAD 525(O2P) | | | |
| | | | | A:LEU 276(N) | 3.04 | B:FAD 525(O2P) | | | |
| | | | | A:SER 277(N) | 3.18 | B:FAD 525(O2P) | | | |
| | | | | A:HIS 375(N) | 3.53 | B:FAD 525(O4') | | | |
| | | | | A:SER 277(OG) | 3.17 | B:FAD 525(O3') | | | |
| | | | | A:HIS 375(ND1) | 3.19 | B:FAD 525(O2') | | | |
| | | | | A:HIS 273(ND1) | 3.12 | B:FAD 525(O1A) | | | |

| | | | | | | |
|---|---|---|---|---|---|---|
| | | | | A:ASN 281(ND2) | 3.13 | B:FAD 525(O2) |
| | | | | A:GLY 274(N) | 2.82 | B:FAD 525(O2A) |
| | | | | A:SER 277(OG) | 2.91 | B:FAD 525(O3B) |
| | | | | A:ASP 406(O) | 3.53 | B:FAD 525(N3) |
| | | | | A:ASN 415 | 2.98 | B:FAD 525(N6A) |
| | 353.9 | 3.5 | PROTEIN+DLZ [ANTENNA CHROMOPHORE] | A:GLY 114(N) | 3.21 | B:DLZ 527(O5') |
| | | | | A:GLY 9(N) | 3.13 | B:DLZ 527(O2) |
| | | | | A:TRP 54(NE1) | 3.72 | B:DLZ 527(O4) |
| | | | | A:SER 36(N) | 3.13 | B:DLZ 527(O4) |
| | | | | A:ASP 10(OD2) | 2.67 | B:DLZ 527(O5') |
| | | | | A:ASP 10(OD1) | 3.06 | B:DLZ 527(O3') |
| | | | | A:VAL 34(O) | 2.78 | B:DLZ 527(N3) |
| | | | OR | | | |
| | 383.5 | 3.6 | PROTEIN+8HDF [ANTENNA CHROMOPHORE] | A:GLY 114[ N ] | 2.03 | A:HDF 602[ O5'] |
| | | | | A:ARG 115[ N | 3.28 | A:HDF 602[ |

| | | | | | | | | | |
|---|---|---|---|---|---|---|---|---|---|
| | | | | ] | | O5'] | | | |
| | | | | A:ARG 119[ NH1] | 2.11 | A:HDF 602[ O4'] | | | |
| | | | | A:ASP 10[ N ] | 3.79 | A:HDF 602[ O2'] | | | |
| | | | | A:GLY 9[ N ] | 3.06 | A:HDF 602[ O2 ] | | | |
| | | | | A:SER 36[ N ] | 3.31 | A:HDF 602[ O4 ] | | | |
| | | | | A:LYS 49[ NZ ] | 2.26 | A:HDF 602[ O8 ] | | | |
| | | | | A:ASP 10[ OD2] | 3.61 | A:HDF 602[ O5'] | | | |
| | | | | A:ASP 10[ OD1] | 2.76 | A:HDF 602[ O2'] | | | |
| | | | | A:VAL 34[ O ] | 3.05 | A:HDF 602[ N3 ] | | | |
| | | | | A:GLY 407[ O ] | 3.75 | A:HDF 602[ O8 ] | | | |
| | | | OR | | | | | | |
| | 328 | 2.3 | PROTEIN+MTHF [ANTENNA CHROMOPHORE] | A:ASP 332[ N ] | 3.86 | X:MHF 527[ O1 ] | | | |
| | | | | A:GLU 117[ N ] | 3.09 | X:MHF 527[ O4 ] | | | |
| | | | | A:ARG 118[ N ] | 3.83 | X:MHF 527[ O4 ] | | | |
| | | | | A:GLY 114[ O ] | 2.55 | X:MHF 527[ NA2] | | | |
| | 173.1 | -20.8 | PROTEIN+SF4 | X | | | A:CYS | 2.2 | B:SF4 |

| | | | | | | | | | |
|---|---|---|---|---|---|---|---|---|---|
| | | | [REDOX CENTRE] | | | | 463[SG ] | | 526[ FE3 ] |
| | | | | | | | A:CYS 450[SG ] | 2.3 | B:SF4 526[ FE1 ] |
| | | | | | | | A:CYS 447[SG ] | 2.2 | B:SF4 526[ FE4 ] |
| | | | | | | | A:CYS 359[SG ] | 2.2 | B:SF4 526[ FE2 ] |
| | 653.8 | -8.8 | PROTEIN+DNA | A:ARG 158[ NH2] | 3.14 | C:DG 7[ O3'] | | | |
| | | | | A:ARG 164[ NH2] | 3.12 | C:DG 7[ OP1] | | | |
| | | | | A:ARG 189[ NH1] | 2.59 | C:DG 7[ OP2] | | | |
| | | | | A:ARG 192[ NH2] | 2.26 | C:DT 8[ O2 ] | | | |
| | | | | A:ARG 158[ NH2] | 2.44 | C:DT 8[ OP2] | | | |
| | | | | A:ARG 192[ NE ] | 2.63 | C:DT 9[ O2 ] | | | |
| | | | | A:TYR 439[ OH ] | 3.52 | C:DT 9[ O2 ] | | | |
| | | | | A:GLN 492[ NE2] | 3.38 | C:DA 12[ OP2] | | | |
| | | | | A:HIS 375[NE2] | 4.01 | C:DT 8[O4] | | | |
| | 283 | -3.1 | PROTEIN+DNA | A:ARG 485 [NH1] | 4.2 | D:DC 10[O2] | | | |
| | | | | A:LYS 491 [NZ] | 3.6 | D:DA 8 [OP1] | | | |

| | | | | | | |
|---|---|---|---|---|---|---|
| | | | | A:TYR 154 [OH] | 3.8 | D:DC 13 [OP1] |
| **Name** | **Interface Area, (Å$^2$)** | **Gain in solvation energy upon interface formation, ΔG (kcal/mol)** | **Molecules forming complexes** | **Hydrogen bonds** | | |
| 3CVY | 739.6 | -11.8 | PROTEIN+DNA | **PROTEIN** | **DISTANCE Å** | **LIGAND** |
| | | | | A:GLN 299(OE1) | 2.94 | C:DT 8(N3) |
| | | | | A:PHE 420(O) | 3.76 | C:DG 10(N2) |
| | | | | A:GLN 299(NE2) | 3.55 | C:DT 8(O2) |
| | | | | A:HIS 365(NE2) | 3.5 | C:DT 8(O4) |
| | | | | A:ARG 421(NH1) | 3.15 | C:DG 9(OP1) |
| | | | | A:HIS 365(NE2) | 2.78 | C:DT 9(O4) |
| | | | | A:ARG 421(NH1) | 3.12 | C:DG 10(OP2) |
| | | | | A:VAL 422(N) | 3.82 | C:DG 10(O4') |
| | | | | A:SER 424(N) | 2.82 | C:DC 11(OP1) |

| | | | | | | |
|---|---|---|---|---|---|---|
| | | | | A:LYS 431(NZ) | 3.55 | C:DC 11(OP2) |
| | 443.7 | -1.1 | | | | |
| | | | PROTEIN+DNA | A:LYS 161(NZ) | 2.85 | D:DT 14(OP1) |
| | | | | A:ARG 502(NH1) | 3.87 | D:DC 11(OP2) |
| | | | | A:ARG 505(NE) | 3.69 | D:DG 12(OP1) |
| | | | | A:ARG 505(NH2) | 3.87 | D:DC 11(OP1) |
| | 580.2 | -6.3 | PROTEIN+FAD [CHROMOPHORE, REDOX CENTRE] | A:ASN 406(OD1) | 3.11 | A:FAD 521(N6A) |
| | | | | A:ASP 397(O) | 2.61 | A:FAD 521(N3) |
| | | | | A:THR 259(N) | 3.06 | A:FAD 521(O1A) |
| | | | | A:THR 259(OG1) | 2.69 | A:FAD 521(O1A) |
| | | | | A:LYS 246(NZ) | 2.76 | A:FAD 521(O2A) |
| | | | | A:LEU 300(N) | 3.74 | A:FAD 521(O4B) |
| | | | | A:ASP 399(N) | 3.04 | A:FAD 521(O4) |
| | | | | A:HIS 365(ND1) | 2.72 | A:FAD 521(O2') |
| | | | | A:TRP 362(NE1) | 3.28 | A:FAD 521(O5') |
| | | | | A:SER 262(N) | 3.05 | A:FAD |

| | | | | | | | | | |
|---|---|---|---|---|---|---|---|---|---|
| | | | | | | 521(O1P) | | | |
| | | | | A:THR 259(OG1) | 3.46 | A:FAD 521(O1P) | | | |
| | | | | A:LEU 261(N) | 2.85 | A:FAD 521(O1P) | | | |
| | | | | A:THR 259(N) | 3.1 | A:FAD 521(O2P) | | | |
| | | | | A:VAL 260(N) | 2.62 | A:FAD 521(O2P) | | | |
| **Name** | **Interface Area, (Å$^2$)** | **Gain in solvation energy upon interface formation, ΔG (kcal/mol)** | **Molecules forming complexes** | **Hydrogen bonds** | | | **Covalent bonds** | | |
| 4DJA | 619.2 | -6.1 | PROTEIN+FAD [CHROMOPHORE, REDOX CENTRE] | PROTEIN | **DISTANCE Å** | **LIGAND** | **PROTEIN** | **DISTANCE Å** | **LIGAND** |
| | | | | A:ASN 406(OD1) | 2.96 | A:FAD 601(N6A) | | | |
| | | | | A:ASP 397(O) | 2.96 | A:FAD 601(N3) | | | |
| | | | | A:ALA 364(O) | 3.54 | A:FAD 601(O4') | | | |
| | | | | A:HIS 265(ND1) | 2.68 | A:FAD 601(O1A) | | | |
| | | | | A:SER 266(N) | 2.81 | A:FAD 601(O2A) | | | |

| | | | | | | |
|---|---|---|---|---|---|---|
| | | | | A:SER 266(OG) | 2.7 | A:FAD 601(O2A) |
| | | | | A:SER 266(OG) | 3.87 | A:FAD 601(O3B) |
| | | | | A:ASN 273(ND2) | 2.86 | A:FAD 601(O2) |
| | | | | A:HIS 366(ND1) | 2.91 | A:FAD 601(O2') |
| | | | | A:HIS 366(N) | 3.46 | A:FAD 601(O4') |
| | | | | A:ARG 369(NH2) | 3.46 | A:FAD 601(O4') |
| | | | | A:SER 266(OG) | 3.45 | A:FAD 601(O1P) |
| | | | | A:LEU 268(N) | 3.15 | A:FAD 601(O1P) |
| | | | | A:SER 269(N) | 3.02 | A:FAD 601(O1P) |
| | | | | A:SER 266(N) | 3.49 | A:FAD 601(O2P) |
| | | | | A:LEU 267(N) | 2.89 | A:FAD 601(O2P) |
| | | | | A:TYR 363(OH) | 2.7 | A:FAD 601(O2P) |
| | 345.2 | 0.8 | PROTEIN+DLZ [ANTENNA CHROMOPHORE] | A:CYS 32(O) | 2.83 | A:DLZ 602(N3) |
| | | | | A:GLU 37(OE1) | 2.7 | A:DLZ 602(O2') |
| | | | | A:GLU 37(O) | 3.89 | A:DLZ 602(O2') |

| | | | | | | | | | |
|---|---|---|---|---|---|---|---|---|---|
| | | | | A:TYR 40(OH) | 3.04 | A:DLZ 602(O3') | | | |
| | | | | A:ASP 10(OD2) | 2.69 | A:DLZ 602(O5') | | | |
| | | | | A:VAL 34(N) | 3.02 | A:DLZ 602(O4) | | | |
| | | | | A:GLY 9(N) | 2.88 | A:DLZ 602(O2) | | | |
| | | | | A:GLY 105(N) | 2.73 | A:DLZ 602(O5') | | | |
| | 182.9 | -21.4 | PROTEIN+SF4 [REDOX CENTRE] | X | | | A:CYS 441[SG ] | 2.3 | A:SF4 603[ FE4] |
| | | | | | | | A:CYS 454[SG ] | 2.3 | A:SF4 603[ FE2] |
| | | | | | | | A:CYS 350[SG ] | 2.3 | A:SF4 603[ FE3] |
| | | | | | | | A:CYS 438[SG ] | 2.3 | A:SF4 603[ FE1] |
| | 602.9 | -10 | PROTEIN+DNA | A:ASN 177[ ND2] | 3.71 | C:DT 8[ O4'] | | | |
| | | | | A:ARG 433[ NH2] | 3.31 | C:DT 9[ O3'] | | | |
| | | | | A:TYR 430[ OH ] | 2.63 | C:DT 9[ O2 ] | | | |
| | | | | A:ARG 476[ NH1] | 2.68 | C:DT 9[ OP2] | | | |
| | | | | A:ARG 433[ NH2] | 2.29 | C:DG 10[ OP1] | | | |
| | | | | A:ARG 433[ | 3.57 | C:DC 11[ | | | |

| | | | | | | | | | |
|---|---|---|---|---|---|---|---|---|---|
| | | | | NH1] | | OP2] | | | |
| | | | | A:ASP 254[ OD2] | 3.50 | C:DT 8[ N3 ] | | | |
| | | | | A:GLN 479[ OE1] | 2.48 | C:DG 10[ N2 ] | | | |
| | | | | A:HIS 366 [NE2] | 4.5 | C:DT 8 [O4] | | | |
| | | | | A:GLN 306 [NE2] | 4.5 | C:DT 8 [O4] | | | |
| | 283.8 | -5.4 | PROTEIN+DNA | A:TRP 176[ NE1] | 3.69 | D:DT 14[ OP1] | | | |
| | | | | A:GLN 184[ NE2] | 2.44 | D:DC 3[ OP2] | | | |
| | | | | A:HIS 475[ NE2] | 3.29 | D:DC 10[ O2 ] | | | |
| **Name** | **Interface Area, (Å$^2$)** | **Gain in solvation energy upon interface formation, ΔG (kcal/mol)** | **Molecules forming complexes** | **Hydrogen bonds** | | | **Covalent bonds** | | |
| 3ZXS | 619.1 | -6.2 | PROTEIN+FAD [CHROMOPHORE, REDOX CENTRE] | PROTEIN | **DISTANCE Å** | **LIGAND** | **PROTEIN** | **DISTANCE Å** | **LIGAND** |
| | | | | A:ASN 402[ OD1] | 3.05 | A:FAD1509[ N1A] | | | |

| | | | | | | |
|---|---|---|---|---|---|---|
| | | | | A:ASP 393[ O ] | 2.97 | A:FAD1509[ N3 ] |
| | | | | A:ALA 360[ O ] | 3.76 | A:FAD1509[ O4'] |
| | | | | A:HIS 261[ ND1] | 2.71 | A:FAD1509[ O1A] |
| | | | | A:ALA 262[ N ] | 2.73 | A:FAD1509[ O2A] |
| | | | | A:ASN 269[ ND2] | 2.82 | A:FAD1509[ O2 ] |
| | | | | A:HIS 362[ ND1] | 2.98 | A:FAD1509[ O2'] |
| | | | | A:HIS 362[ N ] | 3.49 | A:FAD1509[ O4'] |
| | | | | A:ALA 262[ N ] | 3.50 | A:FAD1509[ O1P] |
| | | | | A:LEU 263[ N ] | 2.91 | A:FAD1509[ O1P] |
| | | | | A:TYR 359[ OH ] | 2.57 | A:FAD1509[ O1P] |
| | | | | A:SER 265[ N ] | 3.19 | A:FAD1509[ O2P] |
| | | | | A:LEU 264[ N ] | 2.94 | A:FAD1509[ O2P] |
| | 351.6 | 1.9 | PROTEIN+DLZ [ANTENNA CHROMOPHORE] | A:ALA 32[ O ] | 2.79 | A:DLZ1511[ N3 ] |
| | | | | A:GLU 37[ OE1] | 2.70 | A:DLZ1511[ O2'] |
| | | | | A:TYR 40[ OH ] | 3.67 | A:DLZ1511[ O2'] |

| | | | | | | | | | |
|---|---|---|---|---|---|---|---|---|---|
| | | | | A:ASP 10[ OD1] | 2.71 | A:DLZ1511[ O3'] | | | |
| | | | | A:TYR 40[ OH ] | 2.91 | A:DLZ1511[ O4'] | | | |
| | | | | A:ASP 10[ OD2] | 2.76 | A:DLZ1511[ O5'] | | | |
| | | | | A:VAL 34[ N ] | 3.21 | A:DLZ1511[ O4 ] | | | |
| | | | | A:GLY 9[ N ] | 3.07 | A:DLZ1511[ O2 ] | | | |
| | | | | A:GLY 105[ N ] | 3.23 | A:DLZ1511[ O5'] | | | |
| | 185 | -21.5 | PROTEIN+SF4 [REDOX CENTRE] | X | | | A:CYS 437[SG ] | 2.3 | A:SF4 1510[ FE1] |
| | | | | | | | A:CYS 434[SG ] | 2.2 | A:SF4 1510[ FE4] |
| | | | | | | | A:CYS 450[SG ] | 2.2 | A:SF4 1510[ FE3] |
| | | | | | | | A:CYS 346[SG ] | 2.2 | A:SF4 1510[ FE2] |
| | 834.9 | -6.9 | PROTEIN+DNA | A:ARG 146[ NH1] | 2.11 | C:DG 7[ O3'] | | | |
| | | | | A:HIS 362[ NE2] | 3.24 | C:DT 8[ O4 ] | | | |
| | | | | A:TYR 420[ | 3.24 | C:DT 9[ | | | |

| | | | | | | | | | |
|---|---|---|---|---|---|---|---|---|---|
| | | | | OH ] | | OP2] | | | |
| | | | | A:ARG 429[ NH2] | 3.71 | C:DG 10[ OP1] | | | |
| | | | | A:ARG 179[ NH1] | 3.50 | C:DG 10[ OP2] | | | |
| | | | | A:Gln 302[NE2] | 3.7 | C:DT 8[N3] | | | |
| | 506.3 | -6.2 | PROTEIN+DNA | A:ARG 142[ NH2] | 3.23 | D:DC 13[ OP1] | | | |
| | | | | A:GLN 144[ NE2] | 3.68 | D:DC 11[ O3'] | | | |
| | | | | A:ARG 156[ NH2] | 3.70 | D:DT 14[ OP1] | | | |
| | | | | A:ARG 472[ NH2] | 3.73 | D:DC 11[ O4'] | | | |
| | | | | A:ARG 478[ NE ] | 2.51 | D:DA 8[ OP1] | | | |
| **Name** | **Interface Area, (Å$^2$)** | **Gain in solvation energy upon interface formation, ΔG (kcal/mol)** | **Molecules forming complexes** | **Hydrogen bonds** | | | **Covalent bonds** | | |
| 8IJY | 590.8 | -7.6 | PROTEIN+FAD [CHROMOPHORE, | **PROTEIN** | **DISTANCE Å** | **LIGAND** | **PROTEIN** | **DISTANCE Å** | **LIGAND** |

| | | | | | | |
|---|---|---|---|---|---|---|
| | | | REDOX CENTRE] | A:ASN 402[ OD1] | 2.92 | A:FAD 603[ N6A] |
| | | | | A:ASP 393[ O ] | 3.00 | A:FAD 603[ N3 ] |
| | | | | A:SER 262[ N ] | 3.13 | A:FAD 603[ O1A] |
| | | | | A:SER 262[ OG ] | 2.74 | A:FAD 603[ O1A] |
| | | | | A:HIS 261[ ND1] | 2.82 | A:FAD 603[ O2A] |
| | | | | A:ASN 269[ ND2] | 2.97 | A:FAD 603[ O2 ] |
| | | | | A:HIS 362[ ND1] | 2.89 | A:FAD 603[ O2'] |
| | | | | A:HIS 362[ N ] | 3.37 | A:FAD 603[ O4'] |
| | | | | A:SER 262[ OG ] | 3.38 | A:FAD 603[ O1P] |
| | | | | A:ALA 265[ N ] | 3.28 | A:FAD 603[ O1P] |
| | | | | A:ILE 264[ N ] | 3.33 | A:FAD 603[ O1P] |
| | | | | A:SER 262[ N ] | 3.47 | A:FAD 603[ O2P] |
| | | | | A:TYR 359[ OH ] | 2.74 | A:FAD 603[ O2P] |
| | | | | A:LEU 263[ N ] | 3.01 | A:FAD 603[ O2P] |
| | 379.4 | 3.7 | PROTEIN+8HDF [ANTENNA | A:VAL 34[ O ] | 2.78 | A:HDF 602[ N3 ] |

| | | | | | | | | | |
|---|---|---|---|---|---|---|---|---|---|
| | | | CHROMOPHORE] | A:ASP 10[ OD1] | 2.73 | A:HDF 602[ O2'] | | | |
| | | | | A:ASP 103[ OD2] | 2.76 | A:HDF 602[ O4'] | | | |
| | | | | A:ASP 10[ OD2] | 2.85 | A:HDF 602[ O5'] | | | |
| | | | | A:GLY 9[ N ] | 2.99 | A:HDF 602[ O2 ] | | | |
| | | | | A:TRP 54[ NE1] | 2.92 | A:HDF 602[ O4 ] | | | |
| | | | | A:SER 36[ N ] | 2.98 | A:HDF 602[ O4 ] | | | |
| | | | | A:LYS 49[ NZ ] | 2.67 | A:HDF 602[ O8 ] | | | |
| | | | | A:GLY 9[ N ] | 3.66 | A:HDF 602[ O2'] | | | |
| | | | | A:ASP 10[ N ] | 3.23 | A:HDF 602[ O2'] | | | |
| | | | | A:HIS 39[ NE2] | 2.78 | A:HDF 602[ O3'] | | | |
| | | | | A:ALA 102[ N ] | 3.48 | A:HDF 602[ O5'] | | | |
| | | | | A:ASP 103[ N ] | 3.10 | A:HDF 602[ O5'] | | | |
| | | | | A:VAL 34[ O ] | 2.78 | A:HDF 602[ N3 ] | | | |
| | 172.7 | -21.1 | PROTEIN+SF4 [REDOX CENTRE] | X | | | A:CYS 437[SG ] | 2.2 | A:SF4 601[ FE1] |
| | | | | | | | A:CYS 346[SG ] | 2.2 | A:SF4 601[ FE2] |

| | | | | | | | | | |
|---|---|---|---|---|---|---|---|---|---|
| | | | | | | | A:CYS 434[SG ] | 2.2 | A:SF4 601[ FE4] |
| | | | | | | | A:CYS 450[SG ] | 2.3 | A:SF4 601[ FE3] |
| | 510.7 | -11.7 | PROTEIN+DNA | A:GLN 171[ NE2] | 3.03 | C:DG 7[ OP2] | | | |
| | | | | A:ARG 145[ NH2] | 3.00 | C:DT 8[ OP2] | | | |
| | | | | A:TYR 426[ OH ] | 3.82 | C:DT 9[ O2 ] | | | |
| | | | | A:ARG 472[ NH2] | 3.55 | C:DT 9[ O4 ] | | | |
| | | | | A:TYR 426[ OH ] | 3.49 | C:DT 9[ N3 ] | | | |
| | 184.6 | -1.9 | PROTEIN+DNA | A:ARG 179[ NH2] | 3.21 | D:DA 2[ O5'] | | | |

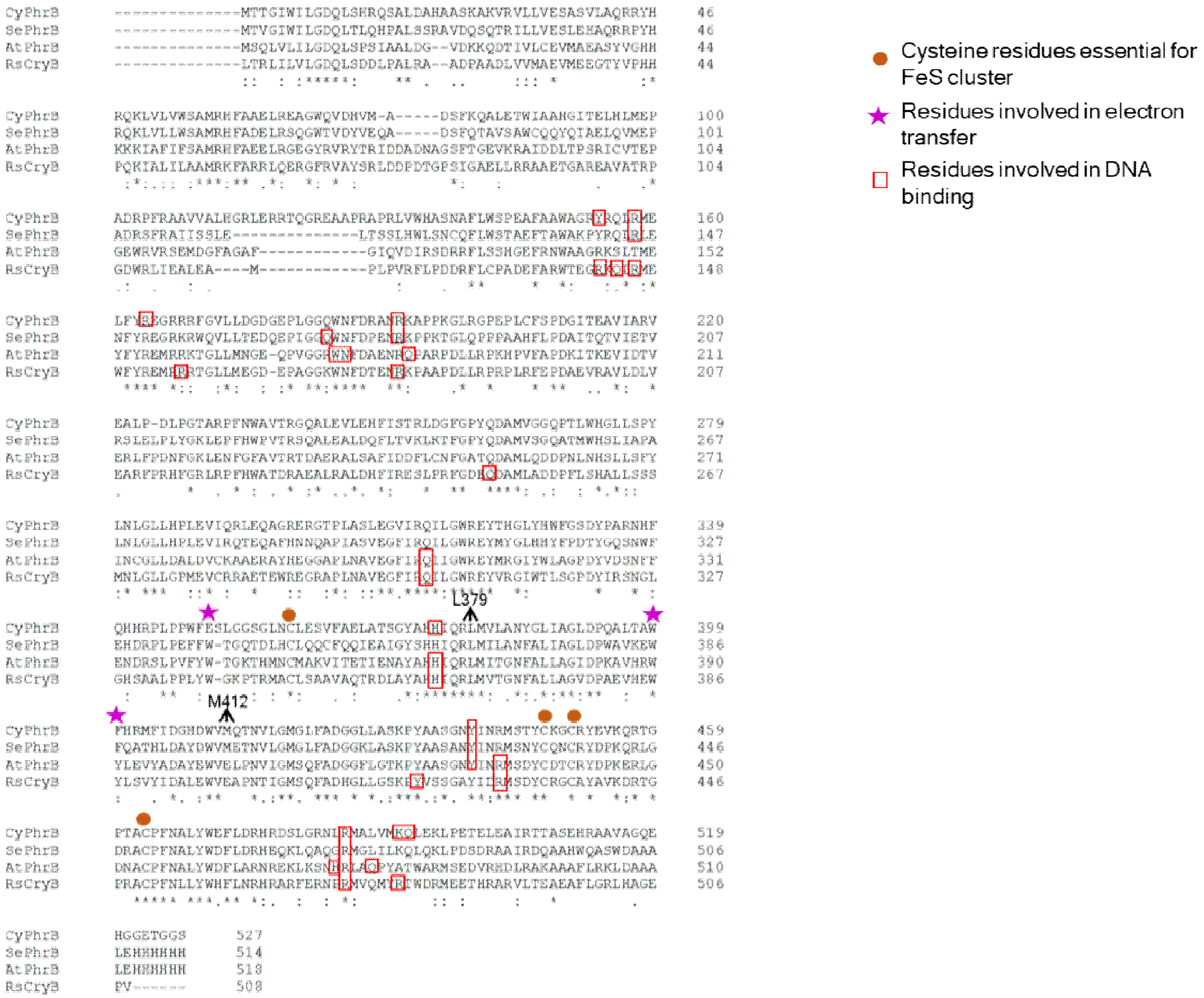
CyPhrB
SePhrB
AtPhrB
RsCryB
L379
M412
Cysteine residues essential for FeS cluster
Residues involved in electron transfer
Residues involved in DNA binding

### Figure S1

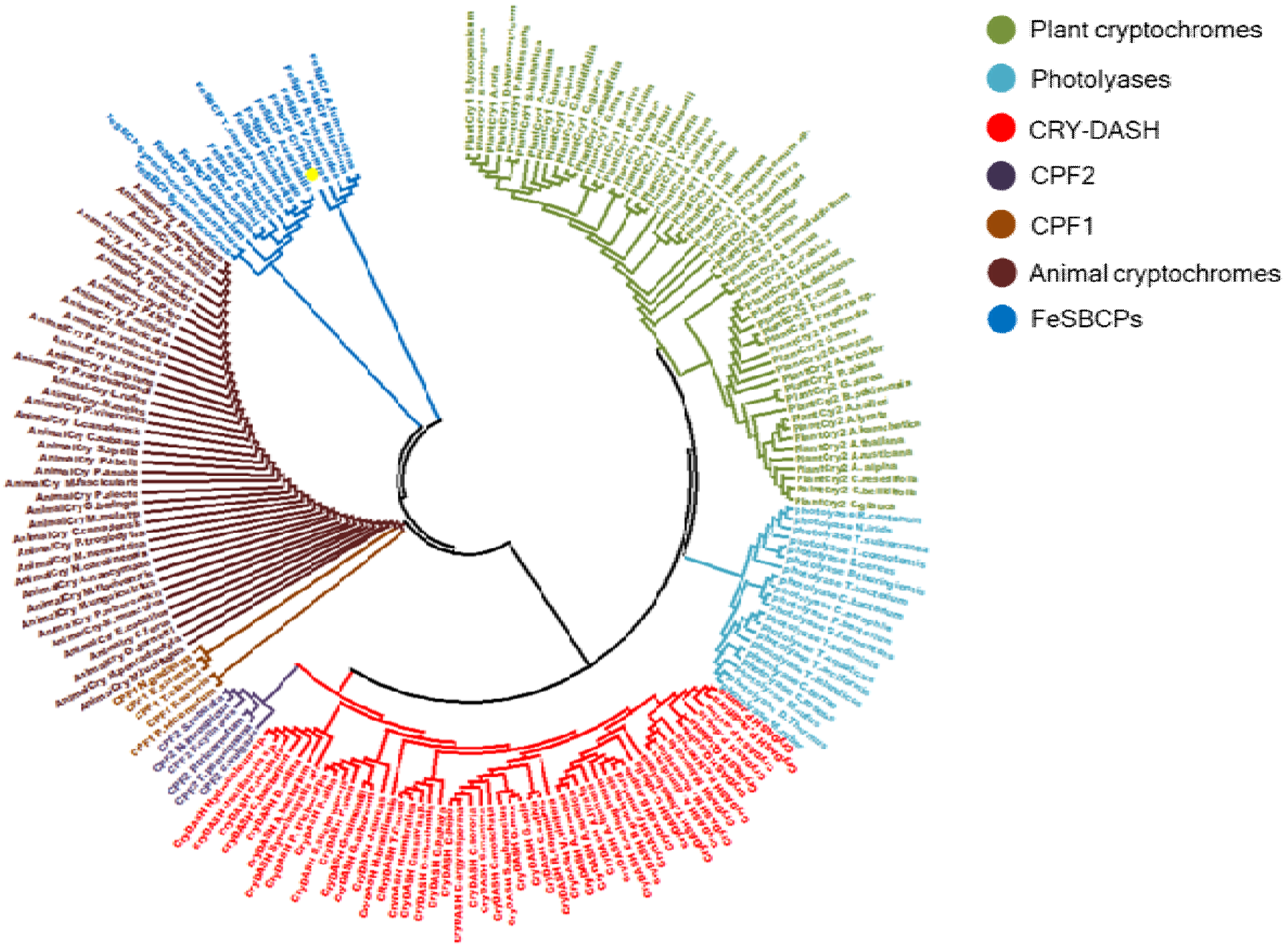
Plant cryptochromes
Photolyases
CRY-DASH
CPF2
CPF1
Animal cryptochromes
FeSBCPs

**Figure S2**

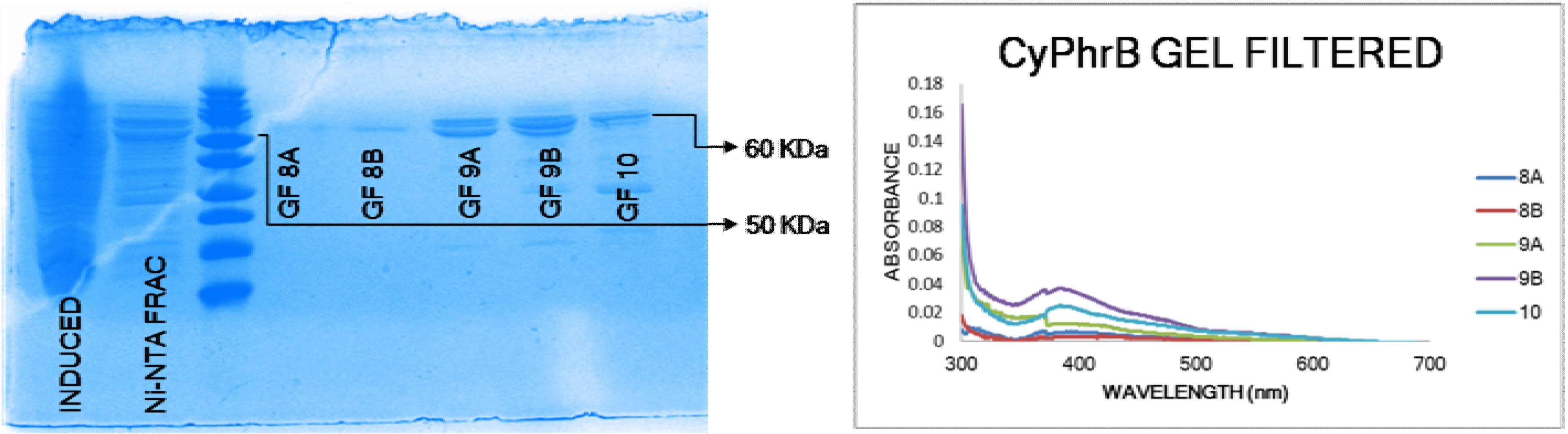
INDUCED
Ni-NTA FRAC
GF 8A
GF 8B
GF 9A
GF 9B
GF 10
60 KDa
50 KDa
CyPhrB GEL FILTERED
0.18
0.16
0.14
0.12
0.1
0.08
0.06
0.04
0.02
0
ABSORBANCE
300
400
500
600
700
WAVELENGTH (nm)
8A
8B
9A
9B
10

**Figure S3**

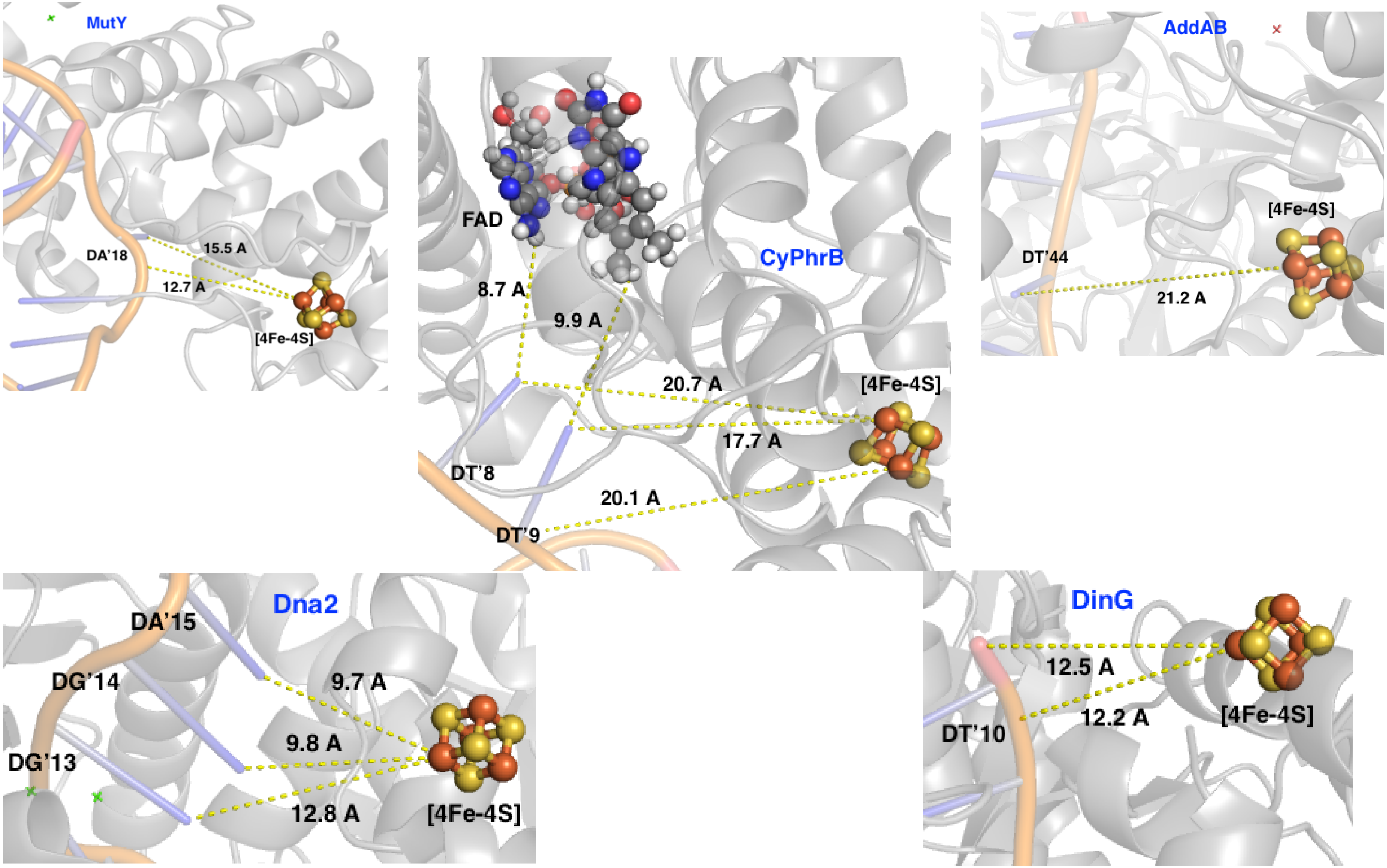
MutY
DA'18
15.5 A
12.7 A
[4Fe-4S]
FAD
8.7 A
9.9 A
CyPhrB
20.7 A
[4Fe-4S]
17.7 A
DT'8
20.1 A
DT'9
AddAB
[4Fe-4S]
DT'44
21.2 A
Dna2
DA'15
DG'14
9.7 A
9.8 A
DG'13
12.8 A
[4Fe-4S]
DinG
12.5 A
12.2 A
DT'10
[4Fe-4S]

**Figure S4**
